# Inhibition of fatty acid oxidation as a new target to treat Primary Amoebic Meningoencephalitis by repurposing two well-known drugs

**DOI:** 10.1101/650325

**Authors:** Maarten J. Sarink, Annelies Verbon, Aloysius G.M. Tielens, Jaap J. van Hellemond

## Abstract

Primary Amoebic Meningoencephalitis (PAM) is a rapidly fatal infection caused by the free-living amoeba *Naegleria fowleri*. The disease mostly affects healthy children and young adults after contaminated water enters the nose, generally during recreational water activities. The amoeba migrate along the olfactory nerve to the brain, resulting in seizures, coma and eventually death. Previous research has shown that *Naegleria gruberi*, a close relative of *N. fowleri*, prefers lipids over glucose as an energy source. Therefore, we tested several inhibitors of fatty acid oxidation alongside the currently used drugs amphotericin B and miltefosine. Our data demonstrate that etomoxir, orlistat, perhexiline, thioridazine and valproic acid inhibited growth of *N. gruberi*. Furthermore, additive effects were seen when drugs were combined. Both thioridazine and valproic acid inhibit in vitro growth of *N. gruberi* in concentrations that can be obtained at the site of infection, which is doubtful with the currently used drugs amphotericin B and miltefosine. Both thioridazine and valproic acid have already been used for other diseases. As the development of new drugs and randomized controlled trials for this rare disease is nearly impossible, repurposing drugs is the most promising way to obtain additional drugs to combat PAM. Thioridazine and valproic acid are available drugs without major side-effects and can, therefore, be used as new complementary options in PAM therapy.

## Introduction

The amoeba *Naegleria fowleri* causes Primary Amoebic Meningoencephalitis (PAM), a rapidly fatal disease of the central nervous system (CNS) (1-3). *N. fowleri* is one of the three most common free-living amoebae that can infect humans, the others being *Acanthamoeba* spp. and *Balamuthia mandrillaris*. These amoebae are ubiquitously present, with *N. fowleri* reported on all continents, except Antarctica (4). *N. fowleri* infections occur mostly in healthy children and young adults during recreational water activities, such as swimming, diving and rafting (5, 6). When water containing *N. fowleri* makes contact with the nasal epithelium, the trophozoite stage of the amoeba can migrate along the olfactory nerve, through the cribriform plate to the olfactory bulb within the CNS (2, 3, 7). Once inside the brain, the trophozoites will cause necrosis and acute inflammation, ultimately leading to death in over 95% of the cases (1, 3). There is concern that global warming and changes in the ecosystems that *N. fowleri* inhabits, may lead to more cases worldwide (8, 9). A wide range of antifungals and antibiotics have been used to treat PAM with varying degrees of effectivity. Most evidence is available for Amphotericin B (AMB) and Miltefosine (MIL), but because of the high mortality rate, more effective drugs are urgently needed (10).

Inhibition of metabolic processes essential to microorganisms is a fruitful strategy for the development of effective drugs (11). Several widely used drugs target the metabolism of the pathogen to exert their killing effect, such as the antimalarials atovaquone and proguanil, and the broad-spectrum antihelminthic and antiprotozoal nitazoxanide (12, 13).

Previous research by our group showed that *N. gruberi*, a close relative to *N. fowleri*, prefers fatty acids as a food source (14). This led us to the hypothesis that inhibiting fatty acid oxidation (FAO) could inhibit growth or even kill the amoeba. We identified several drugs that inhibit fatty acid metabolism in different parts of this pathway. As the metabolic machinery of *N. gruberi* is highly similar to that of *N. fowleri* (14-16), we used *N. gruberi* as a model organism to determine the effects of these compounds. We then compared the effects of these inhibitors to the currently used treatment (AMB and MIL) and determined additive effects of the compounds when they were combined.

## Materials and Methods

### Chemicals and amoeba culture

*N. gruberi* strain NEG-M (ATCC® 30224) was grown axenically at 25 °C in modified PYNFH medium (ATCC medium 1034), as described before (14). Modified PYNFH is composed of peptone, yeast extract, yeast nucleic acid, folic acid, 10% heat-inactivated fetal bovine serum, 100 units/ml penicillin, 100 µg/ml streptomycin and 40 µg/ml gentamicin. All experiments were performed using trophozoites harvested during the logarithmic phase of growth. Amphotericin B (AMB), etomoxir (ETO), miltefosine (MIL), thioridazine (TDZ), orlistat (ORL), perhexiline (PHX) and valproic acid (VPA) were purchased from Sigma. Translucent 96 wells plates were purchased from Greiner Bio-One.

### Growth curves

To determine the effects of fatty acid oxidation inhibitors and current therapies for PAM, 96-well plates were inoculated with 1 × 10^4^ *N. gruberi* trophozoites in PYNFH per well. Compounds were tested per plate in triplicate in at least two independent experiments; controls contained equivalent concentrations of compound solvents (water, PYNFH or DMSO). Optical Density (OD) measurements of the 96 wells were performed every 24 hours using a FLUOstar OPTIMA microplate reader. Regrowth capacity was assessed by collecting contents of the wells at day 5, followed by washing three times with PYNFH to remove the inhibitors, after which the samples were added to a new plate. Controls were diluted 10× to allow proper detection of regrowth in these relatively densely populated amoeba samples.

### Imaging of amoeba

In selected combination experiments, images were taken with an Olympus XY51 phase-contrast microscope on day 0, day 5 (both before and after washing) and on day 14 of experiments and were analysed using Cell^F software (Olympus). To visualize activity of the amoeba, movies were recorded on day 14, with 1 frame every 2 seconds for a total time of 1 minute.

### Data analysis

GraphPad Prism 7 was used to process data. Graphs of separate wells were constructed with OD values on Y-axis, time (days) on X-axis. Area under the curve (AUC) was then calculated by GraphPad Prism 7, where after these were combined into a bar chart.

## Results

Initially, a wide range of concentrations of all investigated compounds was used to estimate the concentration of the compounds at which these inhibit replication of the amoeba. Next, a smaller range of concentrations was used, of which the effect on replication was examined by growth curve analysis through AUC calculation. The results are shown in Figure 1. Addition of VPA resulted in inhibition of growth in a concentration dependent manner (Fig. 1A). Addition of PHX resulted in an inhibition of about 50% in most concentrations (Fig. 1B). ETO addition resulted in clear inhibition at concentrations above 600 µM (Fig. 1C), ORL inhibited circa 50% of growth in all tested concentrations (Fig. 1D), TDZ inhibited growth in a concentration dependent manner (Fig. 1E). AMB was very effective at inhibiting growth, inhibiting circa 75 % at concentrations of 0.2 µM and higher (Fig. 1F). Addition of MIL resulted in inhibition in a concentration dependent manner with efficient inhibition of growth at 80 µM (Fig. 1G). As can be seen in Figure 1 in the bars above the individual graphs, there was a difference in the capacity for regrowth after exposure to the different compounds. Amoeba incubated with all concentrations of VPA and ORL showed regrowth for all concentrations used, while PHX consistently prevented regrowth at concentrations ≥ 90 µM. TDZ showed little regrowth at 40 µM concentrations, while ETO showed little regrowth in concentrations over 600 µM. MIL showed regrowth in concentrations below 80 µM and inconsistently blocked regrowth at concentrations over 80 µM. AMB was most effective in preventing regrowth, always blocking regrowth at concentrations of 0.4 µM or higher.

**Figure 1:**
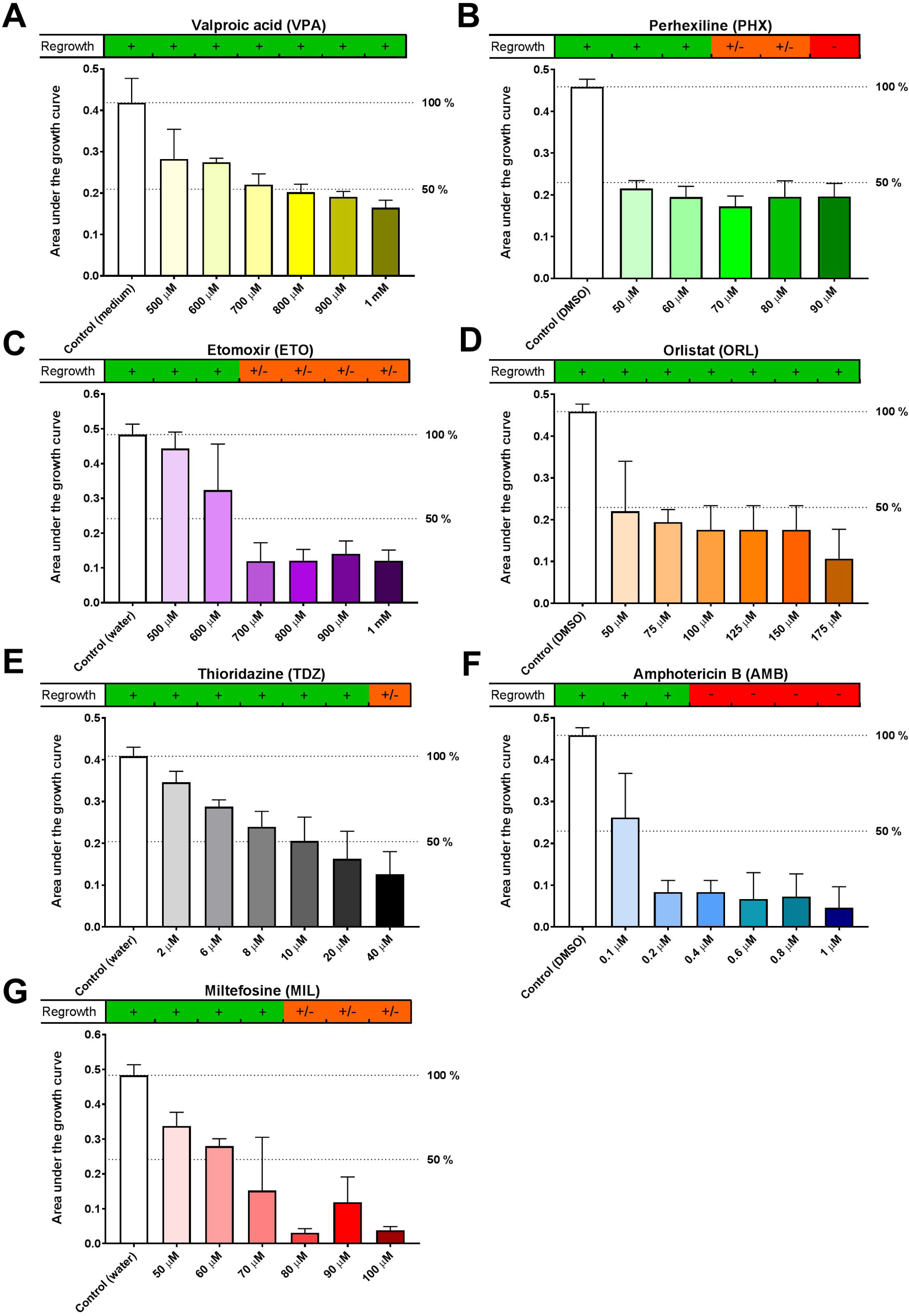
Growth curves of *Naegleria gruberi* were obtained in the presence or absence of inhibitors of fatty acid oxidation or drugs currently used to treat primary amoebic meningitis. Shown is the Area Under the growth Curve (AUC) of compounds and respective controls over a 5-day period. Indicated are lines of 100 % and 50 % of the control AUC. On top of the graph the capacity for regrowth is shown. +: clear regrowth; +/-: inconsistent or little regrowth; -: never any regrowth. Error bars are SD.

Next, compounds were combined to assess whether FAO inhibitors could be a promising addition to the current treatment and to ascertain whether combinations of lower concentrations of FAO inhibitors could enhance the effect of those compounds on their own. For these experiments, VPA, PHX, MIL, ETO, TDZ and ORL concentrations were used which inhibited 50% of growth when used as a single drug. For AMB 0.2 µM was used, as this strongly inhibited growth, but did not kill the amoeba at this concentration. We investigated whether addition of FAO inhibitors could establish killing. Combinations with MIL did not show additive effects when combined with FAO inhibitors or AMB (results not shown). However, an additive effect over the first 5 days was observed when VPA was combined with any of the other FAO inhibitors, as can be seen in Figure 2. VPA alone in a concentration of 700 µM resulted in circa 50 % inhibition of growth, while combining this drug concentration with 50 µM ORL or 60 µM PHX resulted in 75 % inhibition of growth. Combining 700 µM VPA with 600 µM ETO or 10 µM TDZ resulted in an even stronger reduction of growth. However, when the compounds were washed away after 5 days, regrowth was still observed in all combinations. Combinations of the other FAO inhibitors resulted in some, but not evident additive effects on growth during the first 5 days (Fig. S1 A-F).

**Figure 2:**
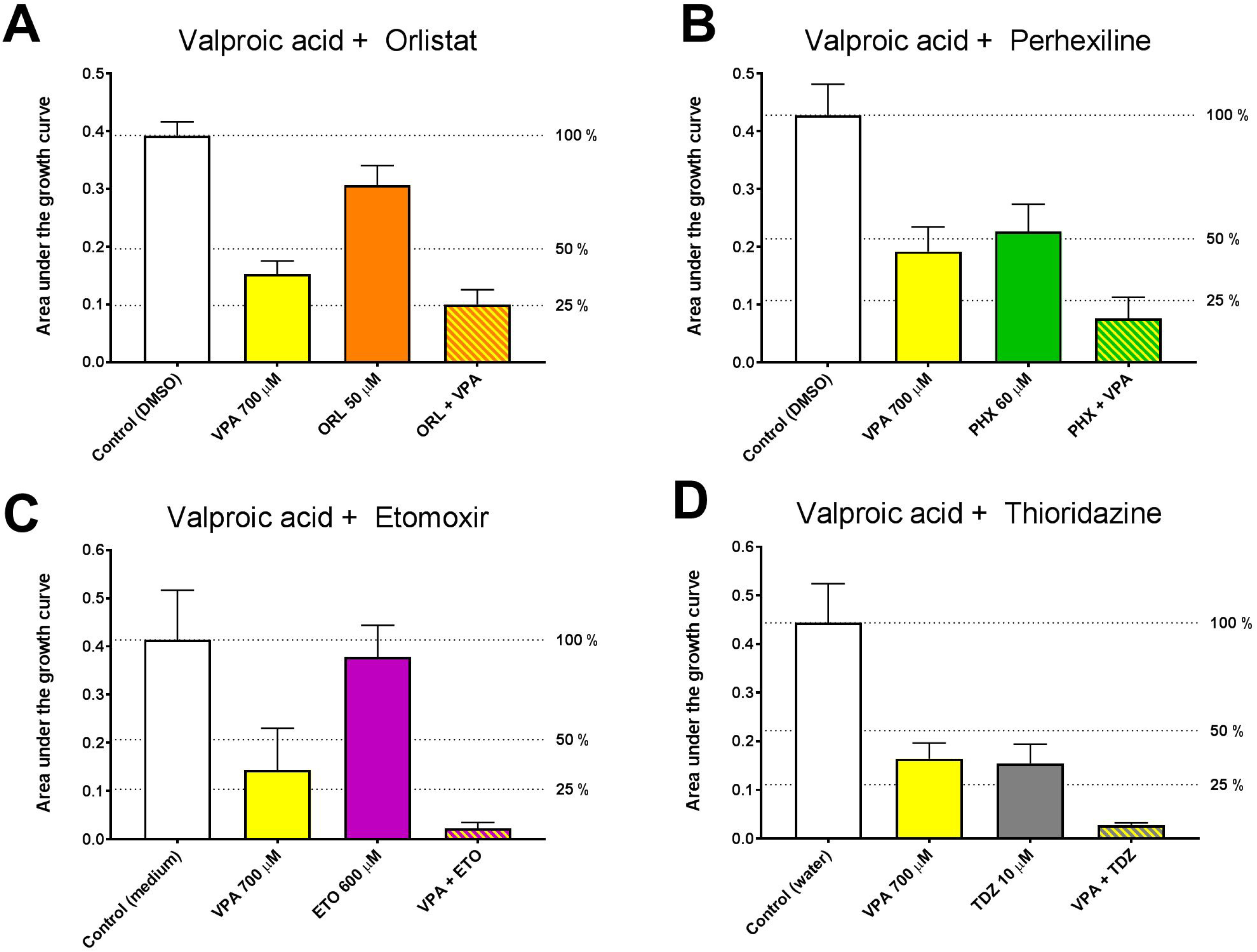
Growth curves of *Naegleria gruberi* were obtained in the presence or absence of valproic acid (VPA), orlistat (ORL), perhexiline (PHX), etomoxir (ETO), thioridazine (TDZ) and combinations of these compounds. Shown is the Area Under the growth Curve (AUC) of compounds and respective controls over a 5-day period. Indicated are lines of 100 %, 50 % and 25 % of the control AUC. Error bars are SD.

Combinations of drugs were also assessed for their capacity to block regrowth. When used separately, none of the compounds used in concentrations in the combination experiments prevented regrowth of the amoeba. However, regrowth was blocked when AMB was combined with ETO or PHX as can be seen in Figure 3 A-B. When AMB was combined with VPA, regrowth capacity was reduced substantially (Fig. 3C). To assess viability of the amoeba, pictures were recorded at day 0, 5 and pictures and movies were recorded at day 14, which is 9 days after removal of the drugs (supplementary material). At day 5 after washing, only active trophozoites were seen in the wells with a single drug exposure (Fig S2 A-D). In contrast, only rounded immobile amoeba were visible in the wells with AMB + ETO and AMB + PHX (Fig. S3 A-B). We noticed one single trophozoite besides the many rounded amoeba in the wells with AMB + VPA (Fig. S3 C). At day 14 the amoeba incubated with AMB + ETO, AMB + PHX or AMB + VPA showed only rounded and disfigured amoeba (Fig. S4 A-C). No visible processes such as movement or vacuolar transport could be detected (See movies S1-4), strongly suggesting that the amoeba that did not regrow were killed by the treatment. In contrast, when compounds were used separately, removal of the inhibitor resulted in dense growth with a mixture of trophozoites and cysts (Fig. S5 A-D).

**Figure 3:**
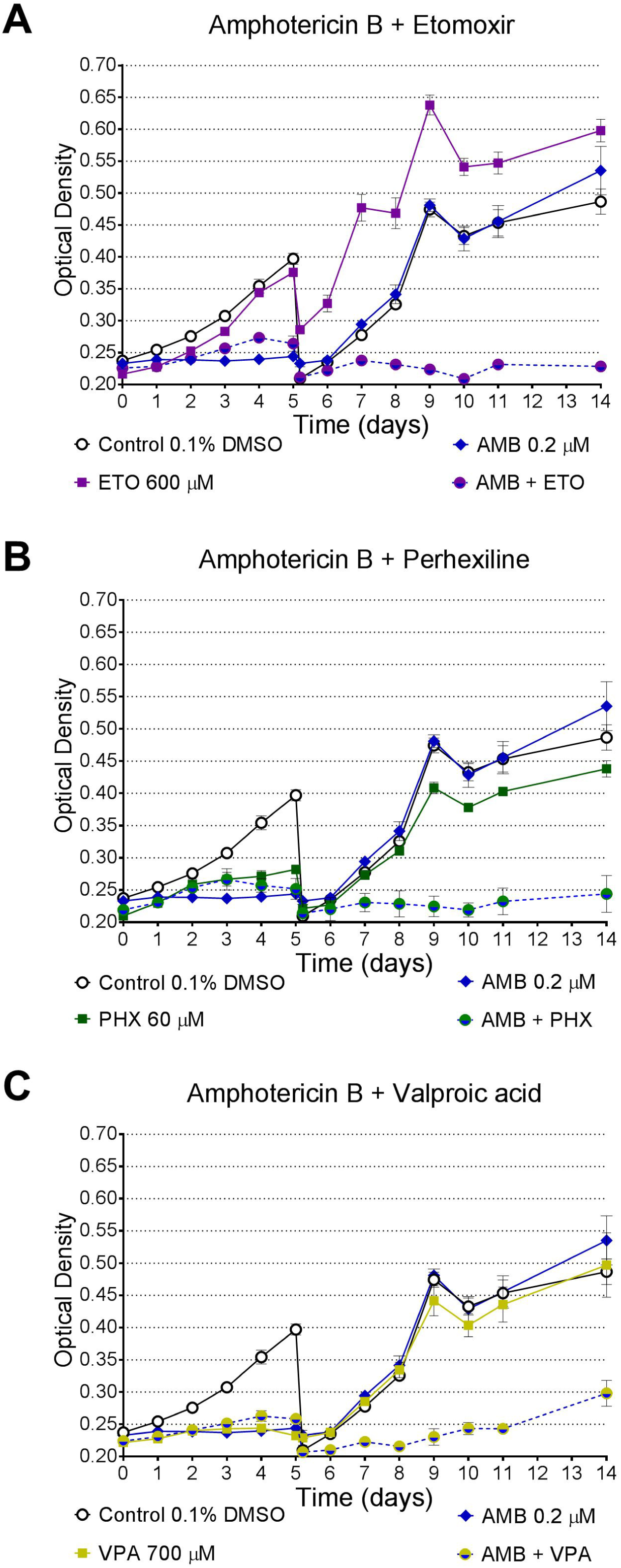
Growth curves of *Naegleria gruberi* in the presence or absence of amphotericin B 0.2 µM (AMB), etomoxir 600 µM (ETO), perhexiline 60 µM (PHX) and valproic acid 700 µM (VPA) and in combinations of these compounds. At day 5, well contents were collected and washed thrice with PYNFH to remove the inhibitors, after which the samples were added to a new plate. The control was diluted 10 times to allow proper detection of regrowth. Shown is a representative example of two independent duplicate experiments, each performed in triplicate.

## Discussion

Our study showed that FAO inhibitors clearly inhibited growth of *N. gruberi in vitro*. Even more, FAO inhibitor PHX killed the amoeba in concentrations ≥ 90 µM. Hence, lipids are not only the preferred food source for *N. gruberi*, but oxidation of fatty acids also seems to be essential for growth. The current treatment of Miltefosine and Amphotericin B was also shown to be effective in inhibiting growth, which is in agreement with previous reports (17-19) and validates our high through-put assay to detect compounds that inhibit growth of *N. gruberi*.

It is notoriously hard to determine whether *Naegleria* cysts are viable or not, because their energy metabolism can be too low to detect, and because their shell is impenetrable for metabolic staining. Therefore, we approached this problem in another way, by giving the amoeba the opportunity to regrow in a nutrient-rich environment (PYNFH) after removal of inhibitory compounds by successive washes. Subsequently, growth of amoeba was followed up to nine days after washing, after which the ability for regrowth was assessed. Of course, it cannot be concluded that amoebae that did not show regrowth were definitely killed, but the chances that amoebae that did not show growth after nine days in such a nutrient-rich environment will be viable and replicate in another environment are very slim.

The investigated FAO inhibitors affect different enzymes involved in lipid metabolism. Thioridazine (TDZ) inhibits peroxisomal oxidation of lipids (20, 21). Orlistat (ORL) inhibits lipases, enzymes that hydrolyse triacylglycerol, thereby obstructing the first step in the breakdown of lipids (22). Etomoxir (ETO) and perhexiline (PHX) inhibit the carnitine palmitoyltransferase-1 (CPT-1), blocking transport of fatty acids into mitochondria (23, 24). Among other activities, valproic acid (VPA) interferes mainly with mitochondrial β-oxidation (25). All these compounds inhibited amoebal growth at various concentrations on their own, but relatively high concentrations were required. However, when compounds were combined, more potent effects were observed. Over the first five days, a clear additive effect was observed when VPA was combined with TDZ, ORL, PHX and ETO. There was also a tendency for enhanced activity when ORL was combined with ETO and TDZ. These results show that when multiple enzymes in lipid catabolism are blocked, this can result in enhanced efficacy of the compounds. Furthermore, when PHX, ETO and VPA were combined with AMB, regrowth of amoeba was prevented, showing that inhibition of the fatty acid oxidation pathway can be a valuable addition to the current treatment regimen.

VPA is one of the oldest anticonvulsants available, present in the WHO list of Essential Medicines and is therefore widely available and used across the world (26, 27). When VPA is prescribed in a conventional dosing regimen in children and adolescents, the maximum concentrations observed in blood can go up to 900 µM without major side-effects (28). We have shown here that VPA can inhibit 50 % growth at concentrations around 700 µM, making it a promising new complementary drug for the treatment of PAM. Possible additional evidence for the efficacy of VPA can be deduced from a described case of a 62-year old male with seizures and a positive PCR result for *N. fowleri* on CSF and brain material. This patient received VPA as a treatment for his seizures and survived, while not receiving any antiparasitic drugs for the amoebal infection (29). TDZ has been in use as an antipsychotic drug since the early 1950s and is now being repurposed as an anti-cancer, anti-inflammatory and antimicrobial agent (30-32). TDZ has been shown to accumulate in brain tissue of chronically treated patients, resulting in concentrations 10-fold higher than that in serum (33). This is in contrast with AMB and MIL, known to be present in very low concentrations in the brain, which could possibly explain the poor prospects for treatment of patients with PAM (34, 35). In a recent clinical study, the sum of TDZ and its metabolites in serum approached 10 µM (32). Since 10 µM TDZ substantially inhibited growth of the amoeba *in vitro*, our results suggest that TDZ is also a promising new drug to treat PAM. Although QTc prolongation can be a side-effect of TDZ, this can be monitored and controlled in a clinical setting. As shown by our results, the addition of both VPA and TDZ resulted in synergistic effects, highlighting the promising nature of combinations of these drugs to treat PAM.

*N. gruberi* is a non-pathogenic relative of *N. fowleri* and genomic analysis has shown that its metabolic machinery is highly similar to that of *N. fowleri* (14-16). Therefore, the observations in our *N. gruberi* model indicate that inhibition of fatty acid oxidation as a new treatment strategy for PAM seems promising. Such a treatment could be directly applied through the repurposing of existing drugs. Development and testing of new drugs for this neglected disease is very difficult, as randomized controlled trials for the treatment of PAM are impossible due to the rapidly fatal nature of the disease and its relatively rare occurrence. It can be argued that if there is evidence that (1) a drug is effective in concentrations attainable in the human body, (2) has few side-effects and (3) is effective against one of the symptoms of PAM, this warrants direct clinical application. Therefore, we propose that if a patient with PAM is experiencing seizures, VPA should be the drug of choice due to the additional inhibiting effects on growth of the amoeba shown in this study. Even more, the combination of the two well-known drugs VPA and TDZ can be a valuable addition to the currently recommended treatment.

